# Evaluation of Phosphogypsum and Pore Volume Water Rates for Reclaiming Saline-Sodic Cambisols of Metehara Sugar Estate, Central Rift Valley of Ethiopia

**DOI:** 10.64898/2026.03.11.710977

**Authors:** Kebenu Feyisa, Kibebew Kibret, Sheleme Beyene, Lemma Wogi

## Abstract

Soil salinity and sodicity are among the major challenges threatening agricultural productivity in the Central Rift Valley of Ethiopia. A column experiment was conducted in laboratory on saline-sodic soils of Metehara Sugar Estate to evaluate the effectiveness of phosphogypsum and leaching in reclaiming these soils. The treatments comprised of five rates of phosphogypsum equivalent to 50, 75, 100, 150, and 200% gypsum requirement, 100% gypsum requirement of natural gypsum, and an absolute control with no amendments applied, and five volumes of leaching water. The treatments were arranged in Complete Randomized Design with three replications. The leaching water was applied to the columns in an intermittent ponding mode. Leachates and soil samples collected from the columns after termination of the leaching process were analyzed for selected soil properties. Results showed that applying phosphogypsum at a rate of 100% gypsum requirement or higher (which is equivalent to ≥ 13 tons/ha) along with 3-4 pore volume of leaching water was found to be the most effective combination to reduce salinity and sodicity to levels that are suitable for most crops (ECe <4 dS/m and ESP < 10%,). The efficiency of phosphogypsum equivalent to 200% gypsum requirement was 81% and 75% in soluble salt removal and Na reduction, respectively. Results of the study suggest that phosphogypsum is a promising reclamation material for saline-sodic soils. However, a field experiment has to be conducted to evaluate the effectiveness of these amendments under natural conditions and come-up with implementable rate recommendations.

## 1. Introduction

Soil degradation, which is driven by human activities [1, 2] and deteriorates soil quality [3], is a critical global issue, with negative consequences on the amount of arable land and food security. Salinity and sodicity, which lead to the formation of salt-affected soils, are among the major forms of chemical soil degradation that pose a serious threat to agricultural productivity globally [3].

Ethiopia, where over 60% of its territory lies in arid and semiarid regions, is among the African countries that are mostly affected by salinity [4]. Empirical evidences indicate that nearly 44 million hectares (36% of the total land area) of the country’s land is potentially vulnerable to salt problems. Out of these 11 million hectares are already affected by varying degrees of salinity and sodicity, with most of these areas concentrated in the Rift Valley [5, 6]. These figures make Ethiopia, the country with the highest percentage of salt-affected land in Africa and the 7^th^ highest globally. As a result, its **i**rrigated lands are threatened by reduced productivity, resulting in increased food insecurity in those regions [7, 6].

For sodic and saline-sodic soils, chemical amendments are inevitable options for replacing exchangeable sodium. These amendments, most of which are typically calcium-bearing materials or acids, aim to replace sodium with calcium and improve soil structure. The chemical amendments should be accompanied by leaching to remove the replaced sodium and excess salts from the soil solution **[8, 9]**. Effectiveness and cost determine the choice of the best chemical amendments **[10]**. In connection with this, several studies [eg., **11, 12,13]** reported the superiority of phosphogypsum, a byproduct of a fertilizer industry, to the commonly used mined gypsum.

Phosphogypsum, formed when phosphoric acid is extracted from phosphate rock using sulfuric acid **[14]** and is primarily composed of calcium sulfate dihydrate (CaSO_4_.2H_2_O), contains impurities like free phosphoric acid, fluorides, and organic matter **[15]**. Phosphogypsum contains nutrient elements such as calcium, phosphorus, and sulfates, along with trace elements, such as silicon and rare earth metals **[16, 17]**. Despite the production of such a huge quantity, only 10-15% of it has been utilized for reclamation and other purposes **[18]**.

Studies [12, 13] have shown the effectiveness of PG in positively affecting soil properties, including reduction of soil pH and exchangeable sodium percentage (ESP), increase crop yields, and growth of beneficial bacteria such as Streptomyces **[19]**.

In addition to its better effectiveness in positively enhancing soil properties than the commonly used mined gypsum, its presence in large quantities and cheaper prices are making PG gain more importance **[20, 8]**. In consent with this, **[21]** reported that the diminishing stock of mined gypsum and a vast pillage of PG in India, necessitated exploring the possibilities of its use as an amendment of sodic soils. Outbakat *et al* **[8]** claims that PG’s higher solubility and richness in electrolytes than mined gypsum makes it a preferred amendment, particularly for sodic and saline-sodic soils.

[8] reported that PG can be used for reclamation of saline-sodic soils with no negative environmental impacts. In addition to the above positive contribution of PG, a recent report advocates for reclassifying PG from a waste product to a valuable resource [22. Phosphogypsum represents an untapped resource with immense potential to drive sustainability in agriculture and other sectors [22].

In Ethiopia’s Awash River Basin, particularly in the Central Rift Valley, agricultural productivity has declined, and more land has been abandoned, especially in areas with long-term irrigation due to salinity problems [4, 23, 6]. Thus, addressing this challenge is a top priority in this part of the country. Availability of natural or mined gypsum in the area is a challenge, and looking for alternative amendments is imperative. Therefore, one potential solution could be using PG, a material that has been tested and found to be effective in reclaiming saline-sodic and sodic soils in other parts of the world like [21] in India; [18] in Russia; [20] in Spain; [24] in Egypt; [25] in Ukraine; [15, 8, 13] in Morocco; [26] in China and [27] in Kazakhstan].

However, PG has never been tested in Ethiopia. Hence, conducting a study to evaluate the effectiveness of PG in combination with leaching for reclaiming saline-sodic soils of the Ethiopian Central Rift Valley is crucially important. Therefore, this study was conducted to evaluate the effects of PG rates on salinity and sodicity-related properties of saline-sodic soils; to develop leaching curves (desalinization and desodification) for saline-sodic soils amended with different PG rates in order to determine its reclamation efficiency; and to determine the amount of water required to complete the reclamation process.

## 2. Materials and Methods

### 2.1. Description of the soil sampling area

The study was carried out using soil samples collected from Metehara Sugar Estate situated in the Fantale district, East Shewa zone of Oromiya Regional State in the Central Rift Valley of Ethiopia in the Middle Awash Basin. The district’s main town, Metehara, is situated at 190 kilometers southeast of Addis Ababa, close to Lake Basaka. Geographically, the district falls within the coordinates of 8.7° to 9.1° North latitude and 39.7° to 40.1° East longitude (Figure 1).

**Figure 1.**
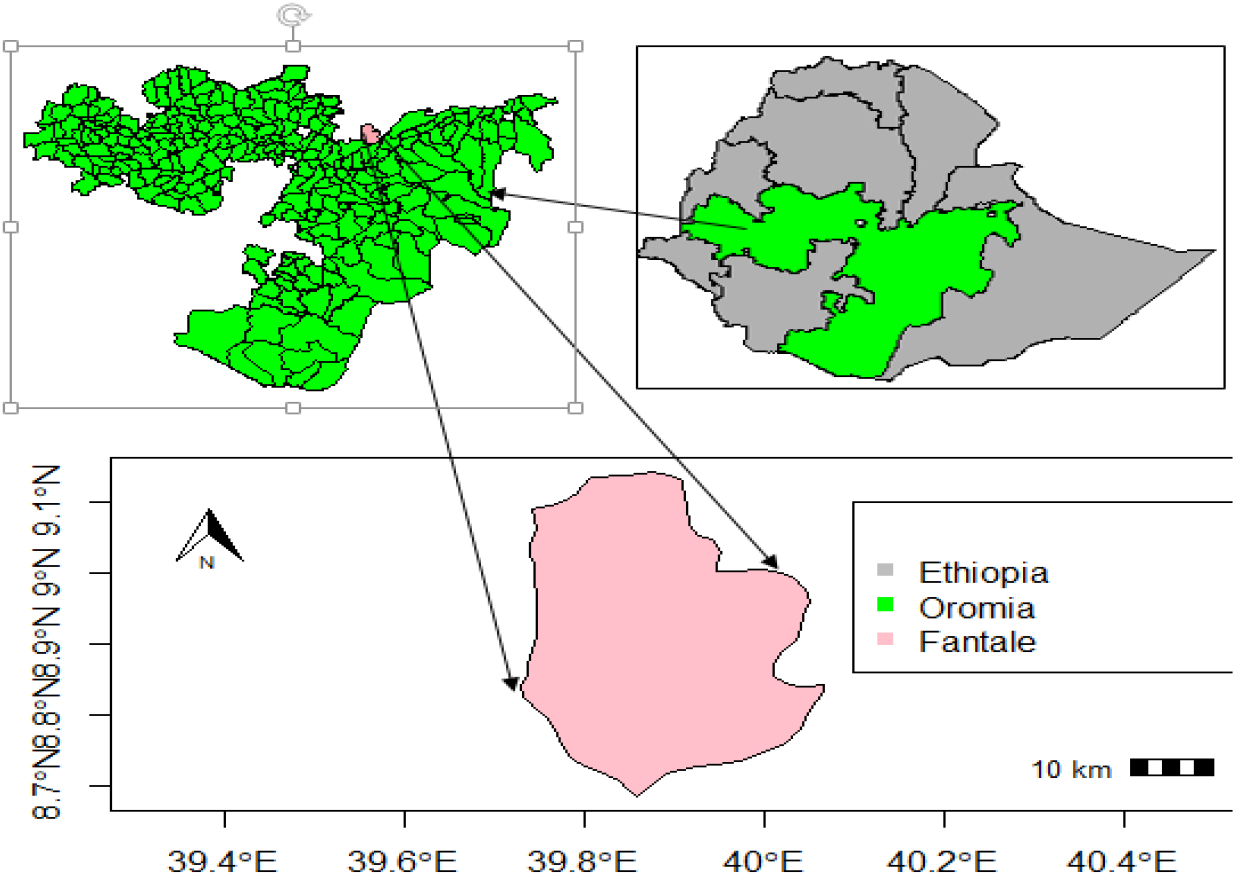
Location map of the soil sampling area.

Since the sugarcane farm was established, Lake Basaka has been expanding significantly, partly due to rising groundwater table and irrigation runoff from nearby canals [28, 29]. This expansion has led to problems like increased salinity in the sugarcane fields, flooding, and disruptions to livestock watering points. The rising water levels have also affected roads and caused concern among local residents of Metehara town [29] and worried the downstream dwellers of Afar Region [28].

#### Climate of the soil sampling area

According to the data obtained from the Fantale District Land Administration and Environmental Protection Office (FDLAEPO, 2009-unpublished) and Ethiopian Meteorological Data Agency (2020-unpublished), the study area has a hot, semi-arid climate with an average annual temperature of 25 °C and about 553 millimeters of annual rainfall. The annual average amount of water that evaporates (1472 mm) is usually more than the rainfall except in July and August (Figure 2). This makes relying on rain for farming unreliable in the area. As a result, irrigation is crucial for agriculture. Due to the dry climate, salts in the soil can’t be washed away by rain.

**Figure 2.**
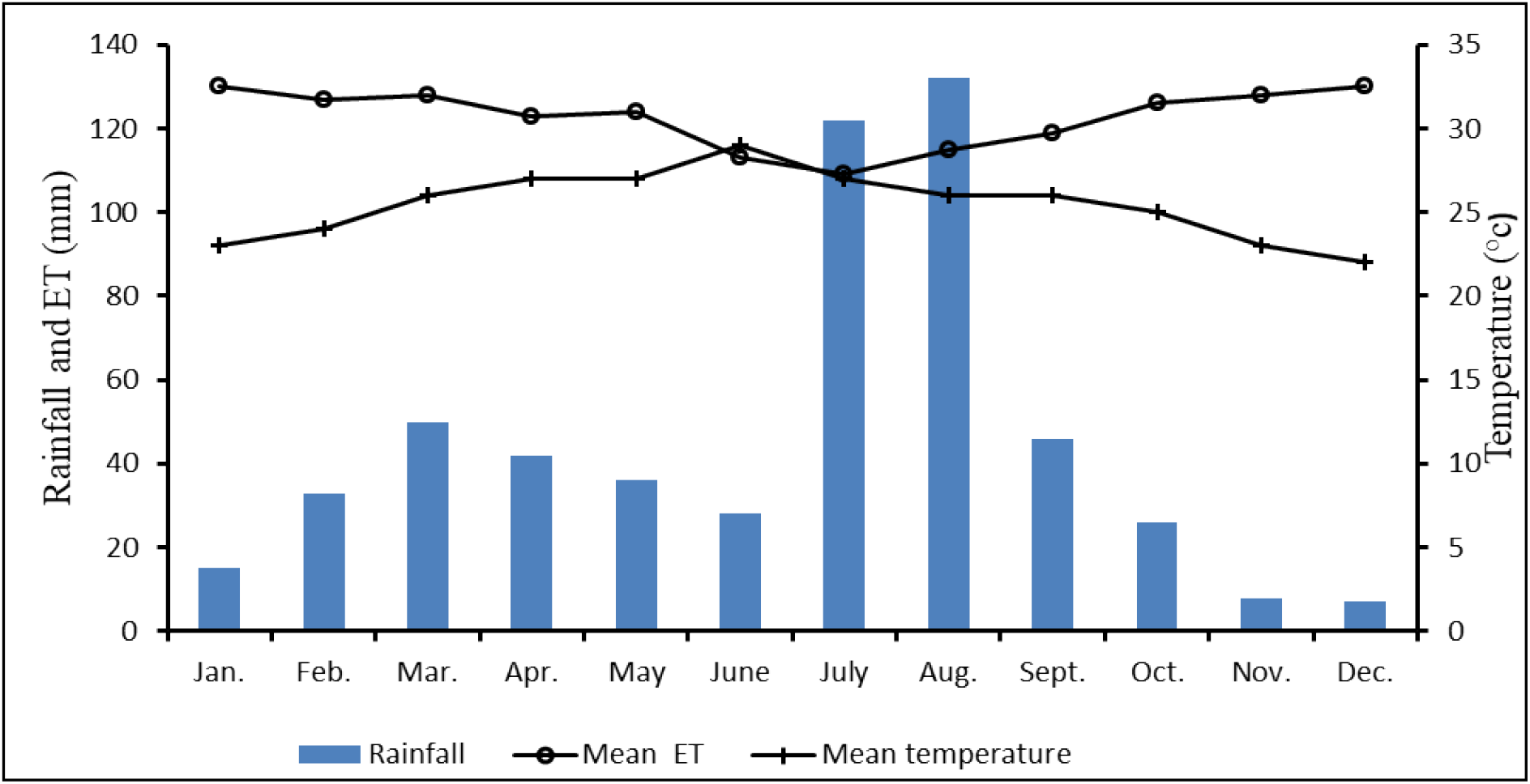
Mean monthly rainfall, ET and temperature of Metehara area for the period 1991-2019 (Source: Ethiopian Meteorological Data Agency, 2020).

#### Geomorphology, Soil and Parent Material of the Soil Sampling Area

The soil sampling area is made up of volcanic rock formations from the Ethiopian Rift System. These rocks are mostly rhyolitic lava, pumice, and tuff. The area is a sunken basin surrounded by hills and mountains. Over time, volcanic material like pumice has been washed into the basin by streams, filling it up. Near the edges of the basin, there are also layers of coarser material [30]

The soils in the study area are mostly volcanic in origin, with some older lake sediments and newer river deposits beside Awash River. These soils are often classified as Calcaric, Vertic, or Hypovertic Cambisols, and they are mostly sandy loam in texture. In parts, especially near the lake, the soils are very salty or alkaline (OWWDSA, 2007-unpublished and [31].

### 2.2. Research Methodology

#### 2.2.1. Soil sample collection and sample preparation

Soil samples used for the column experiment were collected from the top 30 cm depth of salt-affected field of Metehara Sugar Estate using spade. Sub-samples were collected from different points of the field following an X pattern and mixed together thoroughly to make one composite sample. For pre-test analysis of selected soil properties, about 1 kg from the composite sample was taken, air-dried and ground to pass through a 2 mm diameter sieve and taken to the laboratory. The laboratory analyses were carried out in different laboratories depending on availability of facilities. A column leaching experiment was then performed on these samples in laboratory at Haramaya University in 2023.

#### 2.2.2. Laboratory analysis of soil, leaching water, phosphogypsum, and gypsum samples

Particle size distribution was analyzed by the Bouyoucos hydrometer method using sodium hexametaphosphate as a dispersing agent as outlined by Sahlemedhin and Taye [32], while bulk density was determined from undisturbed soil samples following the core method [33]. Finally, total porosity was calculated from the values of bulk density (ρb) and particle density (ρs) with its value of 2.65 g cm^−3^ as:

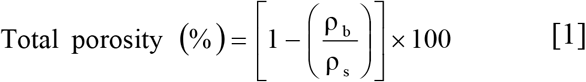

Soil pH was measured using a pH meter based on the potentiometric method outlined by Jackson (1958), using soil-to-water suspensions with a ratio of 1:2.5. The electrical conductivity of the saturated soil paste extracts (ECe) was measured by employing an electrical conductivity meter. To obtain the saturated soil paste, suction was applied using a vacuum pump, as specified by the US Salinity Laboratory Staff [34]. The basic exchangeable cations [calcium (Ca), magnesium (Mg), potassium (K), and sodium (Na)], were extracted from the soil samples by saturating them with a solution of neutral ammonium acetate (1N NH_4_OAc). After the extraction process, the concentrations of exchangeable calcium and magnesium were quantified using an atomic absorption spectrophotometer [35], while the levels of exchangeable sodium and potassium were determined using a flame photometer [36]. The cation exchange capacity (CEC) of the soil was determined by distilling the ammonium-saturated samples using a modified Kjeldahl procedure [37]. Exchangeable sodium percentage (ESP) was calculated from the exchangeable sodium as percent retained by the CEC as:

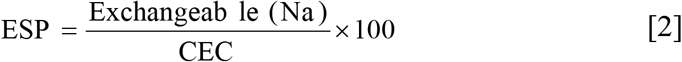

Water soluble basic cations (Ca^2+^, Mg^2+,^ Na^+^ and K^+^) were determined following the procedures described by [38] from the saturated soil paste extracts using atomic absorption spectrophotometer for Ca^2+^ and Mg^2+^ at wavelengths of 422.7 and 285.2 ηm, respectively while K^+^ and Na^+^ were determined using flame photometer at wavelengths of 766.5 and 589.0 ηm, respectively. Carbonate (CO_3_^2-^) and HCO_3_^−^ concentrations were determined from the saturated soil paste extract by simple acidimetric titration using phenolphthalein as an indicator for CO_3_^2-^, and methyl orange for HCO_3_^−^ [38]. Chloride (Cl^−^) was determined by titrating the aliquot used for CO_3_^2-^ and HCO_3_ determinations using silver nitrate to potassium chromate end point while SO_4_^2-^ in the saturated soil paste extracts was determined by precipitating as barium sulphate (BaSO_4_) as described by [38]. According to US salinity laboratory staff [34] the sodium adsorption ratio of the soil solution (SAR) was calculated from the concentrations of soluble Na^+^, Ca^2+^ and Mg^2+^ as :

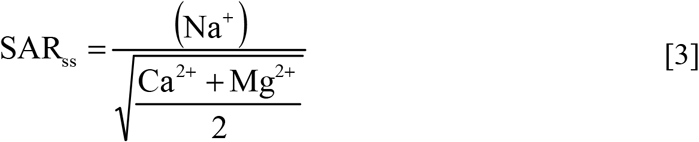

The pH of the irrigation water was assessed using a pH meter, while the electrical conductivity (EC) was measured using an electrical conductivity meter.

#### 2.2.3. Determination of gypsum requirement, treatments and experimental design and procedures

Gypsum requirement (GR) for this particular experiment refers to the amount of gypsum required to reduce the initial exchangeable sodium (30.51%) to a target ESP of 10% for the 30 cm soil depth. The purity level of gypsum used was 88%. The GR was calculated using the following equation [39] while the PG requirements were calculated from the estimated GR:

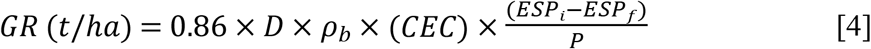

*where D is the depth of the soil to be reclaimed (cm), ρb is the soil bulk-density (g/cm*^*3*^*), CEC is the soil cation exchange capacity (cmol*_*c*_*/kg), ESPi is the initial ESP, ESPf is the target ESP after reclamation and P is the gypsum purity)*.

The treatments comprised of five PG rates (equivalent to 50, 75, 100, 150, and 200% GR), 100% gypsum, and absolute control (no PG or no gypsum application) combined with five pore volumes (PV) of leaching water. The experiment was designed using a completely randomized design (CRD) with three replications, resulting in a total of 21 soil columns. Plastic tubes that are 40 cm tall and 10 cm in diameter were used for the experiment (Figure 3). The bottom of each tube was sealed with a mesh screen and glass wool. A 5 cm sand layer was placed at the bottom of the soil columns to improve water drainage. The columns were filled with 1.1gcm^−3^ bulk density of air-dried soil to a depth of 30 cm and then treated with the different treatment rates. Five pore volumes (PV) of leaching water were then allowed to flow through the columns. The amount of water to be used in leaching was delivered to a saturated soil column in intermittent ponding mode with a water head of 5 cm above the soil surface. The intermittent method is to add equal portions of water to the already saturated soil columns to obtain leachates equal to the added portions and the water that drained out was collected and analyzed. The leaching process continued until the salt concentration in the drained water was equal to the salt concentration of the water used for leaching.

**Figure 3.**
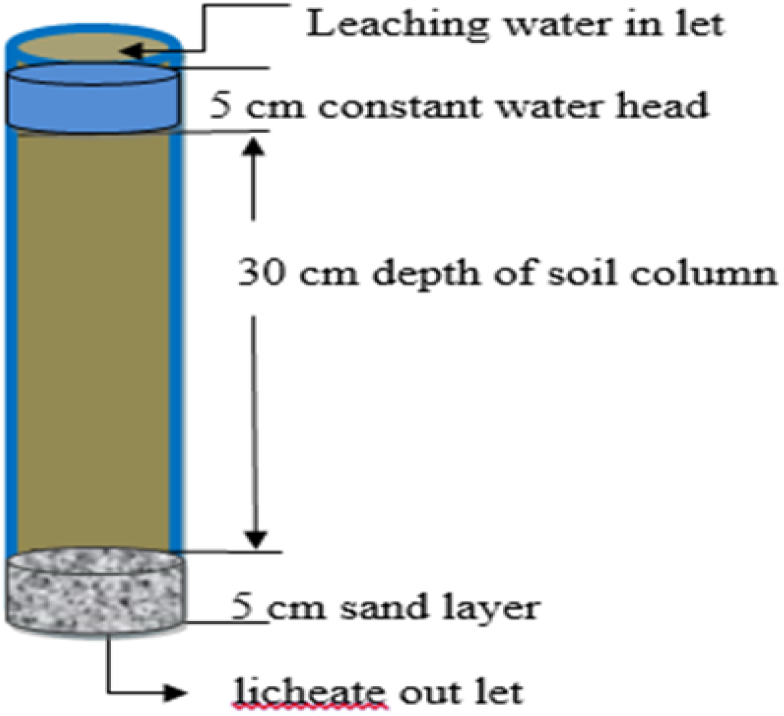
Schematic diagram of the column setup

After mixing amendments with the soil, the soils were leached with tap water having EC of 0.27 dS m^−1^ and SAR of 1.06, indicating that the leaching water was medium in soluble salt concentration (salinity hazard) and low in sodicity hazard according to US Salinity Laboratory Staff, [34] standard. The reclamation requirement (RR), which refers to the amount of water that is needed to reduce the initial soil ECe (salinity) from 5.8 dS m^−1^ to 4.0 dS m^−1^ for 30 cm soil depth, was calculated using the fitted model. Out of the different models tested, the power series model was the best fit for both desalinization and desodification curve. Therefore, to estimate the depth of leaching water for each rate, the following formula was derived from their respective stated equation of the rates.

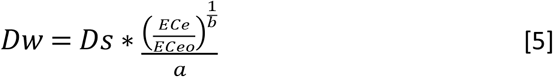

*where a and b are coefficients*

#### 2.2.4. Soil and leachate analysis

After the leaching process was completed, the soil samples were taken out of the tubes, allowed to dry in the air, and then examined for various properties along with the collected leachate. The effects of PG and G rates on soil properties were measured on soil samples collected after the highest (5 PV) leaching water application. The parameters analyzed include electrical conductivity (ECe), pH, cation exchange capacity (CEC), exchangeable sodium percentage (ESP), and exchangeable basic cations. These parameters were analyzed following the procedure described under Section 2.2.2 above.

Furthermore, a leaching curve was created to illustrate the correlation between soil salinity (desalinization) or sodicity (desodification) with the PG rates and depth of water for effective reclamation. This curve was generated by plotting the relative changes in soil salinity and sodicity [40, 41] against the ratio of leaching water to soil depth. The leaching curve serves as a valuable tool for evaluating the effectiveness of the amendments.

The Desalinization Curve was constructed by plotting relative salinity (EC_ef_/ EC_ei_) on the y-coordinate and D_iW_/D_S_ on x-coordinate [41].

*Where;*

EC_ef_ = is soil salinity after an application of specified leaching depth, EC_ei_ = is the initial salinity of the soil, D_iW_ = is the depth of leaching water, D_S_ = is the depth of soil

Similarly, the Desodification curve was constructed by plotting relative changes in soil sodicity by plotting ESP_f_/ ESP_i_ on y-coordinate and D_iW_/D_S_ on the x-coordinate [41].

*Where;*

ESP_f_ = is soil sodicity after an application of specified leaching depth, ESP_i_ = is the initial sodicity of the soil.

#### 2.2.5. Evaluating the efficiency of PG rates and PV of leaching water on soil reclamation

To evaluate the efficiency of the PG rates used as a reclamation material, the changes in ECe and ESP were measured as primary indicators. In this regard, the efficiency of the PG treatment rates for the studied saline-sodic soil was calculated according to [42] at the end of the leaching experiment as follows:

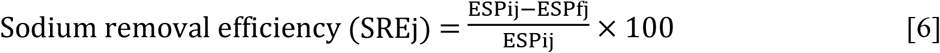

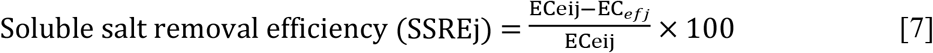

To evaluate the efficiency of combined use of PG and PV of leaching water, an empirical constant (K) was measured and used as another indicator. The K value for the studied saline-sodic soils was calculated according to [41]:

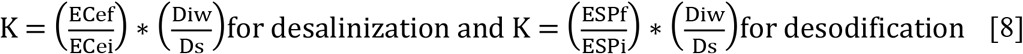

*where;* K is an empirical coefficient

ECef = soil salinity after an application of specified leaching depth, ECei = the initial salinity of soil, Di_W_ = the depth of leaching water, D_S_ = the depth of soil, ESPf = exchangeable sodium after an application of specified leaching depth, ESPi = the initial exchangeable sodium of soil, j=the j^th^ rate of PG or PV of leaching water

#### 2.2.6. Statistical Analysis

A two-way analysis of variance (ANOVA) was conducted using R Statistical software to assess the impact of applied phosphogypsum rates and leaching water on saline-sodic soil and its residual. In instances where significant differences were observed, Tukey’s test (P<0.05) was employed to separate the means, allowing for the determination of the most suitable desodification and desalinization curves for the tested soils.

## 3. Results and Discussion

Based on the analyzed data (Table 1), the textural class of the analyzed soil was dominantly sandy loam while its pH showed alkaline reaction. Using the salt affected soil classification criteria set by [42], the soil of the study field is classified as saline-sodic, i.e., it has both soluble salt and high sodium content that satisfy the criteria for saline-sodic soil category. This indicates that the experimental soil requires an amendment for normal plant growth and production. The PG used was acidic, while the purity of the gypsum was 88%. According to [34] rating standard, the leaching water used was medium in its salinity and low in sodicity hazards (Table 1).

**Table 1.**
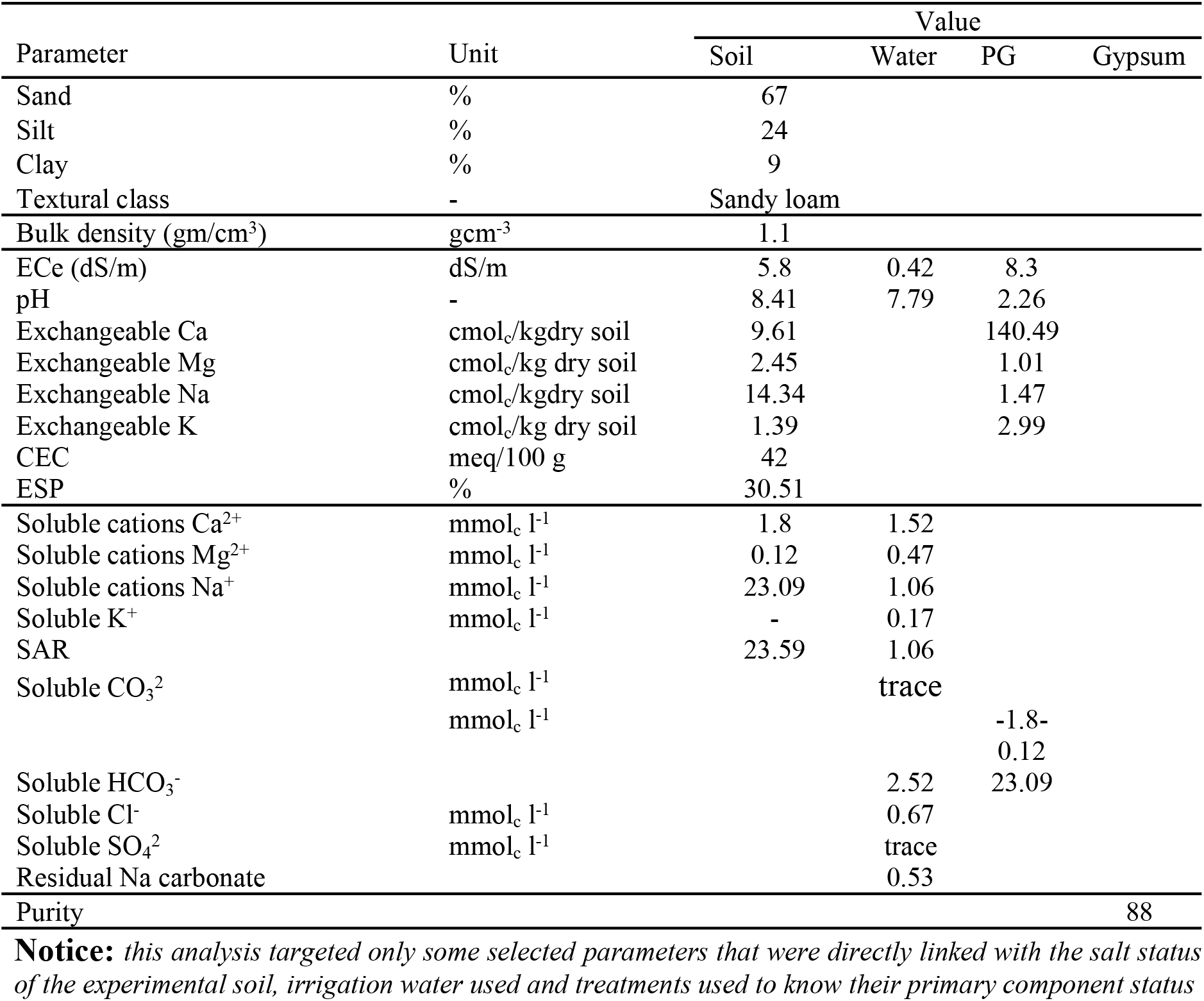
Some properties of the experimental soil (0-30 cm), irrigation water, phosphogypsum, and gypsum used for this study.

### 3.1. Leachate Analysis

#### 3.1.1. Effect of PG and PV water rates on pH, EC and SAR

The interaction effect of PG and PV leaching water on soil pH, EC, soluble cations, and SAR was significant (P ≤ 0.05) (Table 2). Accordingly, the lowest pH (7.5) was recorded from the combined use of PG at 200% GR and 5 PV leaching water, and 100% gypsum and 5 PV leaching water (Table 2). On the other hand, the highest pH (8.7) was obtained from treatments that involved zero PG application and 3 and 4 PV leaching water, after most water was leached through the soil (Table 2). The decrease in pH might be associated with the acidic nature of PG which is expected to reduce the pH and the releasing of calcium ions, which replace sodium ions from the soil’s exchange complex. The removal of the replaced sodium ions from the soil solution might have been facilitated by the applied leaching water at the highest pore volume level. In line with the current finding, similar studies [11, 43, 26] conducted on saline-sodic soils’ reclamation using PG reported that certain acidic components in PG can neutralize bicarbonate ions in the soils, and further contribute to the decrease in soil pH.

**Table 2.** Interaction effect of PG X PV leaching water on pH, EC, SAR and soluble cations (Ca^2+^, Mg^2+^, Na^+^, K^+^) of the leachates from a column experiment.

| PG rates | PV | pH | EC<br>(dS/m) | Soluble cations (meq/l) |  |  |  | SAR |
| --- | --- | --- | --- | --- | --- | --- | --- | --- |
|  |  |  |  | Ca | Mg | Na | K |  |
| 0%GR | 1 | 8.3<sup>cdef</sup> | 10.6<sup>a</sup> | 1.8<sup>q</sup> | 0.12<sup>i</sup> | 23.1<sup>f</sup> | 0.1<sup>c</sup> | 23.3<sup>c</sup> |
|  | 2 | 8.5<sup>abc</sup> | 7.5<sup>d</sup> | 1.8<sup>q</sup> | 0.13<sup>i</sup> | 23.2<sup>f</sup> | 0.1<sup>c</sup> | 23.4<sup>c</sup> |
|  | 3 | 8.7<sup>a</sup> | 7.6<sup>d</sup> | 1.7<sup>q</sup> | 0.13<sup>i</sup> | 23.5<sup>ef</sup> | 0.1<sup>c</sup> | 24.8<sup>b</sup> |
|  | 4 | 8.7<sup>a</sup> | 7.6<sup>d</sup> | 1.5<sup>q</sup> | 0.12<sup>i</sup> | 23.6<sup>ef</sup> | 0.2<sup>c</sup> | 26.2<sup>a</sup> |
|  | 5 | 8.4<sup>bcd</sup> | 7.5<sup>d</sup> | 1.5<sup>q</sup> | 0.12<sup>i</sup> | 23.9<sup>e</sup> | 0.1<sup>c</sup> | 26.6<sup>a</sup> |
| 50%GR | 1 | 8.3<sup>cdef</sup> | 10.6<sup>a</sup> | 9.2<sup>p</sup> | 0.7<sup>h</sup> | 20.9<sup>g</sup> | 0.3<sup>abc</sup> | 9.3<sup>gh</sup> |
|  | 2 | 8.2<sup>cdefg</sup> | 6.0<sup>f</sup> | 10.5<sup>lm</sup> | 1.0<sup>defgh</sup> | 19.2<sup>i</sup> | 0.1<sup>c</sup> | 8.0<sup>ij</sup> |
|  | 3 | 8.1<sup>defghi</sup> | 5.7<sup>f</sup> | 10.0<sup>mn</sup> | 1.0<sup>defgh</sup> | 17.0<sup>k</sup> | 0.1<sup>c</sup> | 7.2<sup>kl</sup> |
|  | 4 | 8.2<sup>defgh</sup> | 6.0<sup>f</sup> | 11.3<sup>ij</sup> | 1.1<sup>cdefg</sup> | 15.8<sup>l</sup> | 0.1<sup>c</sup> | 6.3<sup>lm</sup> |
|  | 5 | 8.0<sup>efghi</sup> | 5.7<sup>f</sup> | 12.5<sup>fg</sup> | 0.9<sup>gh</sup> | 14.5<sup>m</sup> | 0.2<sup>bc</sup> | 5.6<sup>mno</sup> |
| 75%GR | 1 | 8.4<sup>bcd</sup> | 10.6<sup>a</sup> | 9.4<sup>op</sup> | 0.9<sup>gh</sup> | 25.5<sup>d</sup> | 0.5<sup>a</sup> | 10.7<sup>f</sup> |
|  | 2 | 8.3<sup>cdef</sup> | 8.5<sup>c</sup> | 9.9<sup>no</sup> | 1.8<sup>a</sup> | 20.1<sup>h</sup> | 0.3<sup>abc</sup> | 8.3<sup>hi</sup> |
|  | 3 | 8.2<sup>cdefg</sup> | 6.0<sup>f</sup> | 10.7<sup>kl</sup> | 1.5<sup>b</sup> | 16.0<sup>l</sup> | 0.2<sup>bc</sup> | 6.8<sup>kli</sup> |
|  | 4 | 8.1<sup>defghi</sup> | 6.8<sup>e</sup> | 12.1<sup>gh</sup> | 1.0<sup>defgh</sup> | 13.3<sup>n</sup> | 0.2<sup>bc</sup> | 5.2<sup>nop</sup> |
|  | 5 | 8.0<sup>efghij</sup> | 5.1<sup>g</sup> | 12.7<sup>f</sup> | 0.9<sup>gh</sup> | 11.2<sup>p</sup> | 0.1<sup>c</sup> | 4.3<sup>pq</sup> |
| 100%GR | 1 | 8.2<sup>cdefg</sup> | 10.1<sup>ab</sup> | 10.5<sup>lm</sup> | 0.9<sup>gh</sup> | 26.5<sup>c</sup> | 0.2<sup>bc</sup> | 11.1<sup>ef</sup> |
|  | 2 | 8.2<sup>cdefg</sup> | 5.9<sup>f</sup> | 12.1<sup>gh</sup> | 1.3<sup>bcd</sup> | 20.5<sup>gh</sup> | 0.5<sup>a</sup> | 7.9<sup>ijk</sup> |
|  | 3 | 7.9<sup>ghij</sup> | 4.6<sup>gh</sup> | 13.5<sup>cd</sup> | 1.3<sup>bcd</sup> | 16.8<sup>k</sup> | 0.2<sup>bc</sup> | 6.2<sup>lmn</sup> |
|  | 4 | 7.8<sup>ijk</sup> | 3.2<sup>i</sup> | 12.9<sup>ef</sup> | 0.9<sup>gh</sup> | 13.4<sup>n</sup> | 0.1<sup>c</sup> | 5.1<sup>nop</sup> |
|  | 5 | 7.7<sup>jk</sup> | 3.1<sup>i</sup> | 14.3<sup>ab</sup> | 1.0<sup>defgh</sup> | 12.5<sup>o</sup> | 0.2<sup>bc</sup> | 1.1<sup>s</sup> |
| 150%GR | 1 | 8.2<sup>cdefg</sup> | 9.7<sup>b</sup> | 11.3<sup>ij</sup> | 1.1<sup>cdefg</sup> | 27.9<sup>b</sup> | 0.2<sup>bc</sup> | 11.9<sup>e</sup> |
|  | 2 | 8.3<sup>cdef</sup> | 4.6<sup>gh</sup> | 12.1<sup>gh</sup> | 1.2<sup>bcd</sup> | 18.2<sup>j</sup> | 0.2<sup>bc</sup> | 7.1<sup>kl</sup> |
|  | 3 | 7.9<sup>ghij</sup> | 4.5<sup>h</sup> | 13.8<sup>bcd</sup> | 1.8<sup>a</sup> | 13.6<sup>n</sup> | 0.5<sup>a</sup> | 4.9<sup>op</sup> |
|  | 4 | 7.8<sup>ijk</sup> | 2.6<sup>ij</sup> | 14.1<sup>ab</sup> | 1.4<sup>bc</sup> | 10.1<sup>q</sup> | 0.2<sup>bc</sup> | 3.6<sup>q</sup> |
|  | 5 | 7.7<sup>jk</sup> | 2.1<sup>j</sup> | 14.6<sup>a</sup> | 1.2<sup>bcd</sup> | 7.4<sup>r</sup> | 0.1<sup>c</sup> | 1.0<sup>s</sup> |
| 200%GR | 1 | 8.3<sup>cdef</sup> | 10.6<sup>a</sup> | 11.8<sup>hi</sup> | 0.9<sup>gh</sup> | 33.3<sup>a</sup> | 0.2<sup>bc</sup> | 13.2<sup>d</sup> |
|  | 2 | 8.2<sup>cdefg</sup> | 4.2<sup>h</sup> | 12.8<sup>f</sup> | 0.9<sup>gh</sup> | 16.2<sup>l</sup> | 0.2<sup>bc</sup> | 6.4<sup>lm</sup> |
|  | 3 | 7.9<sup>ghij</sup> | 3.8<sup>hi</sup> | 13.9<sup>bc</sup> | 1.1<sup>cdefg</sup> | 12.6<sup>o</sup> | 0.2<sup>bc</sup> | 4.6<sup>opq</sup> |
|  | 4 | 7.7<sup>jk</sup> | 2.6<sup>ij</sup> | 14.0<sup>bc</sup> | 1.2<sup>bcd</sup> | 6.5<sup>s</sup> | 0.4<sup>ab</sup> | 2.4<sup>r</sup> |
|  | 5 | 7.5<sup>k</sup> | 2.0<sup>j</sup> | 14.6<sup>a</sup> | 1.3<sup>bcd</sup> | 5.1<sup>t</sup> | 0.3<sup>abc</sup> | 0.9<sup>s</sup> |
| 100%<br>gypsum | 1 | 8.2<sup>cdefg</sup> | 10.1<sup>ab</sup> | 11.0<sup>jk</sup> | 1.0<sup>defgh</sup> | 25.1<sup>d</sup> | 0.3<sup>abc</sup> | 10.2<sup>fg</sup> |
|  | 2 | 8.2<sup>cdefg</sup> | 6.6<sup>e</sup> | 11.9<sup>h</sup> | 1.2<sup>bcd</sup> | 20.9<sup>g</sup> | 0.2<sup>bc</sup> | 8.2<sup>hi</sup> |
|  | 3 | 7.9<sup>ghij</sup> | 5.0<sup>g</sup> | 13.3<sup>de</sup> | 1.3<sup>bcd</sup> | 17.7<sup>jk</sup> | 0.3<sup>abc</sup> | 6.5<sup>lm</sup> |
|  | 4 | 7.7<sup>jk</sup> | 4.6<sup>gh</sup> | 14.0<sup>bc</sup> | 0.9<sup>gh</sup> | 14.5<sup>m</sup> | 0.5<sup>a</sup> | 5.3<sup>nop</sup> |
|  | 5 | 7.5<sup>k</sup> | 3.6<sup>hi</sup> | 14.3<sup>ab</sup> | 1.0<sup>defgh</sup> | 7.6<sup>r</sup> | 0.2<sup>bc</sup> | 2.8<sup>r</sup> |
| PG rates |  | *** | *** | *** | *** | *** | * | *** |
| PV |  | *** | *** | *** | *** | *** | ns | *** |
| PG rates*PV |  | ** | *** | *** | *** | *** | ** | *** |
| CV (%) |  | 2.3 | 5.1 | 3.0 | 20.4 | 1.9 | 7.4 | 7.8 |
| LSD<sub>0.05</sub> |  | 0.3 | 0.5 | 0.5 | 0.3 | 0.5 | 0.2 | 1.1 |

\*\* and \*\*\*, means significant at $p < 0.01$ and $p < 0.001$ respectively, while 'ns' means not significant at $p < 0.05$ . CV is the coefficient of variation. Means followed by the same letter(s) within a column are not significantly different at $p \leq 0.05$ . LSD is the least significant difference at $p \leq 0.05$ , PG is phosphogypsum as treatment while PV is the pore volume of leaching water.

Similarly, combined use of PG at the highest rate (200% GR) and 5PV leaching water resulted in the lowest EC (2 dS/m) value as compared with the other treatments, while the highest EC value (≥ 10.0 dS/m) was recorded from the control and all the PG treatments when used in combination with 1 PV leaching water (Table 2). This implies that combined use of PG and leaching water at the highest rates was the most effective treatment in leaching salts out of the soil column. The results further showed that lowering the soil salinity to a safe level (< 4 dS/m) can be achieved from combined application of PG at the rates of 100% GR and 150% GR with ≥ 4 PV, 200% GR with ≥ 3 PV, and 100% gypsum with 5 PV leaching water (Table 2). Furthermore, these results revealed that reducing the salinity to a safe level is possible by combining lower rates of PG with decreasing amount of leaching water while much more water needed with 100% pure gypsum. This in turn shows PG is more effective than pure gypsum in minimizing soil salinity in the study area.

The decrease in salinity is likely due to the leaching of salt and the positive impact of PG on soil structure as it releases calcium that replaces sodium from the exchange site. In line with the current findings, several other similar studies [44, 45, 13] confirmed the ability of PG in improving soil structure and reducing the negative effects of excess sodium. Moreover, these studies demonstrated that applying PG to saline-sodic soils reduces salinity with the application of less leaching water. Furthermore, they ascribed the increase in salt concentration in the water that drains from the soil to the release of calcium and sulphate ions from PG.

The sodium adsorption ratio (SAR) in the leachate was also highly significantly (P < 0.001) influenced by the interaction effects of PG and PV leaching water, with the highest reduction in SAR occurring with the combined use of the highest rates of PG and PV leaching water (Table 2). However, applying PG at a rate of ≥50% GR in combination with ≥ 1 PV leaching water was enough to reduce the sodicity to the safe (< SAR value of 13) level (Table 2).

Similarly, application of PG at100% GR in combination with 1 or more PV leaching water was able to reduce the SAR value to less than the safe value. The reduction of the SAR level to values less than 13 with application of 50% GR followed by leaching with just 1 PV leaching water indicates the effectiveness of PG in reclaiming saline-sodic soils. Owing to the good internal drainage of the soil (sandy loam texture) and the easily soluble nature of PG in water, less than 5 pore volumes of leaching water was enough to achieve a safe level of sodium content for most crops (SAR < 13). In consent with this, [25] reported that application of 6 t/ha PG to saline-sodic soils in combination with irrigation water reduced SAR by 81% compared to the control.

#### 3.1.2. Effect of PG and PV water rates on soluble cations (Ca^2+^, Mg^2+^, Na^+^, K^+^)

The interaction effect of phosphogypsum (PG) and PV leaching water on soluble Ca^2+^ was highly significant (P<0.001), with the highest concentration (14.6 meq L^−1^) observed at 150% and 200% GR combined with 5 PV of leaching water (Table 2). The lowest concentration of soluble Ca^2+^ was recorded in the control treatment under all the PV leaching water (Table 2). On the other hand, treatment combinations of 100% GR with 5 PV leaching water, 150% GR with ≥ 4 PV leaching water, 200% GR with 5 PV leaching water, and 100% gypsum requirement with 5 PV leaching water were statistically at par. Furthermore, the soluble Ca^2+^ increased almost consistently with an increase in PG rate as well as PV leaching water, which could be due to the release of Ca^2+^ ions contained in the PG following its dissolution. A study by [45] also reported a similar finding in which increase in the amount of applied PG resulted in an increase in the concentration of Ca^2+^ in the soil solution, which could easily be leached by the applied water.

Unlike calcium, sodium (Na^+^) levels in the soil solution were highly significantly (P<0.001) decreased with increasing PG application rate and PV of leaching water (Table 2). All the PG treatments removed most of the sodium (Na^+^) ions from the soil solution during the first PV of leaching water. The highest removal of Na^+^ ions (33.3 meq L^−1^) was observed at the interaction effect of the highest PG application rate (200%GR) and the first PV of leaching water followed by 150% GR when compared to the control and 100% gypsum. This shows that PG was very effective in removing the exchangeable sodium, especially during the first two rounds of leaching which was even more effective than using gypsum. After the initial removal, the amount of sodium remaining in the soil continued to decrease with higher rates of PG applications, as it releases more calcium and replaces the sodium from the exchange sites. A study by [8] found similar results, confirming the PG’s effectiveness in removing sodium from saline-sodic soils. Similarly [45] found that increasing the amount of PG increased the concentration of calcium in the soil and replaced sodium from the exchange site which can then be removed through leaching.

Although the interaction effects of PG and PV leaching water on soluble Mg^2+^ and K^+^ were significant (P < 0.001 and P <0.01, respectively), their concentration was extremely low (Table 2). This is consistent with the findings of other studies, which also found that PG has little to no effect on the concentrations of soluble Mg^2+^ and K^+^ in the soil solution. For example, a study by [46] found that PG had little to no effect on the concentrations of soluble Mg^2+^ and K^+^ ions in the soil solution. The authors attributed this to the fact that PG contains less amount of these cations, and therefore, it does not release a significant amount of Mg^2+^ or K^+^ ions into the soil solution.

### 3.2. Effect of PG and PV of Leaching Water on Selected Soil Properties

#### 3.2.1. Soil pH, ECe and ESP

Results of analysis of variance showed that soil pH, electrical conductivity (ECe), and exchangeable sodium percentage (ESP) were all highly significantly (P < 0.001) affected by the PG rates (Table 3). Accordingly, the lowest soil pH (7.3) was recorded after 5 PV leaching in the soils treated by PG at 200% GR, while the highest pH value (8.9) was recorded in the control treatment which was statistically at par with the 50% GR PG rate. The PG rates equivalent to ≥ 100 GR resulted in pH values that are statistically at par. Application of 200% GR PG accompanied by leaching using 5 PV water decreased the pH from strongly alkaline (> 8.4) to slightly alkaline (7.3). On the contrary, the pH remained in the strongly alkaline range in the soils that were leached with no PG amendment and treated with PG rate equivalent to 50% GR, suggesting that in saline-sodic soils leaching alone and/or application of low rate of amendments is not enough to remove the excess exchangeable Na. In line with the findings of this study, [8, 46] reported that PG was effective in reducing pH, ECe, and ESP of saline-sodic soils, with the highest reductions in these parameters occurring in the soils that received the highest PG application rate (30 t ha^−1^).

**Table 3.** The main effects of PG rates on selected chemical properties of saline-sodic soils from Metehara Sugar Estate, CRV of Ethiopia.

| PG rates | pH | ECe<br>(dS/m) | Exchangeable cations (meq/100 g of soil) |  |  |  | ESP<br>(%) |
| --- | --- | --- | --- | --- | --- | --- | --- |
|  |  |  | Ca | Mg | Na | K |  |
| 0%GR | 8.9<sup>a</sup> | 4.58<sup>a</sup> | 8.6<sup>g</sup> | 1.1<sup>c</sup> | 11.4<sup>a</sup> | 0.9<sup>a</sup> | 31.9<sup>a</sup> |
| 50%GR | 8.8<sup>a</sup> | 4.45<sup>a</sup> | 28.7<sup>f</sup> | 1.9<sup>b</sup> | 10.7<sup>ab</sup> | 0.6<sup>b</sup> | 26.1<sup>b</sup> |
| 75%GR | 8.4<sup>b</sup> | 4.42<sup>a</sup> | 34.3<sup>e</sup> | 1.8<sup>b</sup> | 10.3<sup>b</sup> | 0.5<sup>c</sup> | 22.8<sup>c</sup> |
| 100%GR | 7.4<sup>cd</sup> | 3.84<sup>ab</sup> | 39.7<sup>d</sup> | 3.4<sup>a</sup> | 4.5<sup>c</sup> | 0.19<sup>de</sup> | 9.4<sup>de</sup> |
| 150%GR | 7.4<sup>cd</sup> | 3.26<sup>cd</sup> | 48.1<sup>b</sup> | 3.7<sup>a</sup> | 4.4<sup>cd</sup> | 0.2<sup>d</sup> | 8.0<sup>e</sup> |
| 200%GR | 7.3<sup>dc</sup> | 3.23<sup>c</sup> | 52.9<sup>a</sup> | 4.0<sup>a</sup> | 4.0<sup>d</sup> | 0.1<sup>e</sup> | 7.0<sup>ef</sup> |
| 100%G | 7.6<sup>c</sup> | 3.93<sup>b</sup> | 40.4<sup>c</sup> | 3.5<sup>a</sup> | 4.9<sup>d</sup> | 0.2<sup>d</sup> | 10.0<sup>d</sup> |
| PG rates | *** | *** | *** | *** | *** | *** | *** |
| CV | 1.5 | 7.9 | 2.6 | 12.6 | 5.9 | 10.8 | 3.8 |
| LSD<sub>0.05</sub> | 0.2 | 0.4 | 1.8 | 0.62 | 0.7 | 0.07 | 1.1 |

\*\*\* means strongly significant at 0.001. The same letters within a column indicate no significant difference at $p \leq 0.05$ , CV is the coefficient of variation. LSD is the least significant difference at $p < 0.05$ , PG is phosphogypsum as treatment while G is gypsum.

Similar to the trends observed for pH, the electrical conductivity (ECe) of the soil after leaching also decreased with increasing PG application rate (Table 3). The lowest ECe values (3.26 and 3.23 dS/m) was observed at the higher PG application rates (150% and 200%GR respectively) after leaching with 5 PV of water. This suggests that most of the soluble salts were removed from the soil, and a safe salinity level (<4 dS/m) for most crops was achieved at these PG application rates. Similarly, a study by [11] found that PG was effective in reducing the ECe of saline-sodic soils. The authors found that the PG application rate had a significant effect on reduction in ECe and higher reductions were observed at higher PG application rates.

Similarly, the lowest ESP (7.1%) was recorded after leaching the soil with 5 PV water in the soils treated with PG at a rate equivalent to 200% GR, while the highest (31.9%) was observed in the control treatment (Table 3). Furthermore, the values recorded after treating the soils with 100% pure gypsum, and PG rates at 100% GR and 150% GR were statistically at par. The results further indicated that application of PG at rates equivalent to ≥ 100% GR followed by 5 PV leaching water were able to reduce the ESP level to the target (safe) level of less than 10%. Thus, application of PG at rates ≥100% GR followed by leaching with 5 PV water reduces the ESP of the soils to safe level than applying pure gypsum at 100% GR rat combined with same amount of leaching water. In agreement with the current findings, [12] reported that PG was more effective than natural gypsum in reducing the ESP of saline-sodic soils.

#### 3.2.2. Exchangeable cations (Ca, Mg, Na, K)

The effect of the treatments on all the exchangeable bases was highly significant (P < 0.001) (Table 3). The effects were particularly considerable on exchangeable Ca, Mg, and Na. The highest exchangeable Ca (52.9 cmol/kg soil) was recorded in the soils that were treated with PG at a rate equivalent to 200% GR, whereas the lowest (8.6 cmol/kg soil) was registered in the control treatment (Table 3). Moreover, within the PG rates, Ca cation increased significantly with increase in PG rates, indicating the strong contribution of PG to soil.

With regard to exchangeable Mg, application of PG at a rate equivalent to ≥100% GR and 100% natural gypsum resulted in equal mean values, with the lowest value registered in the soils that were not treated with PG or natural gypsum but only leaching (Table 3). The effect of PG treatments and natural gypsum on exchangeable Nawas also highly significant, with values decreasing from 11.4 cmol/kg in the control treatment to 4.0 cmol/kg in the soil treated with PG at 200% GR rate. Similarly, application of PG at rates equivalent to ≥150% GR and 100% natural gypsum resulted in values of exchangeable Na that are statistically at par.

In line with the current findings, [8, 46] observed similar trends in that application of PG increased the exchangeable calcium content and decreased the exchangeable sodium content of saline-sodic soils. Similarly, [45] indicated that the increase in exchangeable Ca and Mg following the application of PG is associated with the richness of PG in these cations and their subsequent release following its dissolution when applied to the soil. [27] associated the increase in the divalent cations with PG’s contribution to improving soil structure, which likely enhances soil nutritional regimes while reducing the proportion of sodium absorbed.

There are a few dissimilar findings in the literature regarding the effects of phosphogypsum (PG) on the exchangeable cations in the soil. For example, a study by [47] indicated that PG application had little to no effect on exchangeable magnesium (Mg) content of saline and sodic soils. This is in contrast to the findings in Table 3, which showed that PG application significantly increased the exchangeable Mg content of the soil. Others reported no change in case of sodic soil [46] or even a decrease in exchangeable K. This is likely due to the fact that PG is a variable material, and its composition can vary depending on the source and processing method.

### 3.3 Soil Desalinization and Desodification curves

Out of the different models (linear, logarithmic, quadratic and power series) tested, the power series model was the best fit for both desalinization and desodification curves of this research data. The equations for the power series model for each treatment rate are shown in Table 4.

**Table 4.** Relationship between ECef/ECei and Dw/Ds for desalinization and relation between ESPf/ESPi and Dw/Ds for desodification.

| Treatment rates | Relationship between $EC_{ef}/EC_{ei}$ and $D_w/D_s$<br>(Desalinization) | Relationship between $ESP_f/ESP_i$ and $D_w/D_s$<br>(Desodification) |
| --- | --- | --- |
| 0%GR | $Y=0.698x^{-0.211}$, $R^2=0.631$ | $Y=0.875x^{-0.073}$, $R^2=0.307$ |
| 50%GR | $Y=0.537x^{-0.397}$, $R^2=0.702$ | $Y=0.661x^{-0.277}$, $R^2=0.774$ |
| 75%GR | $Y=0.56x^{-0.381}$, $R^2=0.764$ | $Y=0.608x^{-0.339}$, $R^2=0.843$ |
| 100%GR | $Y=0.337x^{-0.809}$, $R^2=0.948$ | $Y=0.469x^{-0.55}$, $R^2=0.927$ |
| 150%GR | $Y=0.251x^{-0.994}$, $R^2=0.913$ | $Y=0.241x^{-1.13}$, $R^2=0.986$ |
| 200%GR | $Y=0.184x^{-1.38}$, $R^2=0.94$ | $Y=0.184x^{-1.38}$, $R^2=0.94$ |
| 100% Gypsum | $Y=0.235x^{-1.08}$, $R^2=0.951$ | $Y=0.251x^{-1.06}$, $R^2=0.912$ |

Where: $Y$ represents $E_{Ce}/E_{Ci}$ (relative salinity) for desalinization and $ESP/ESP_i$ (relative sodicity) for desodification, $x$ represents $Dw/Ds$

The desalinization and desodification curves (Figures 4 and 5, respectively) were established for the tested saline-sodic soil using the developed equations (Table 4).

**Figure 4.**
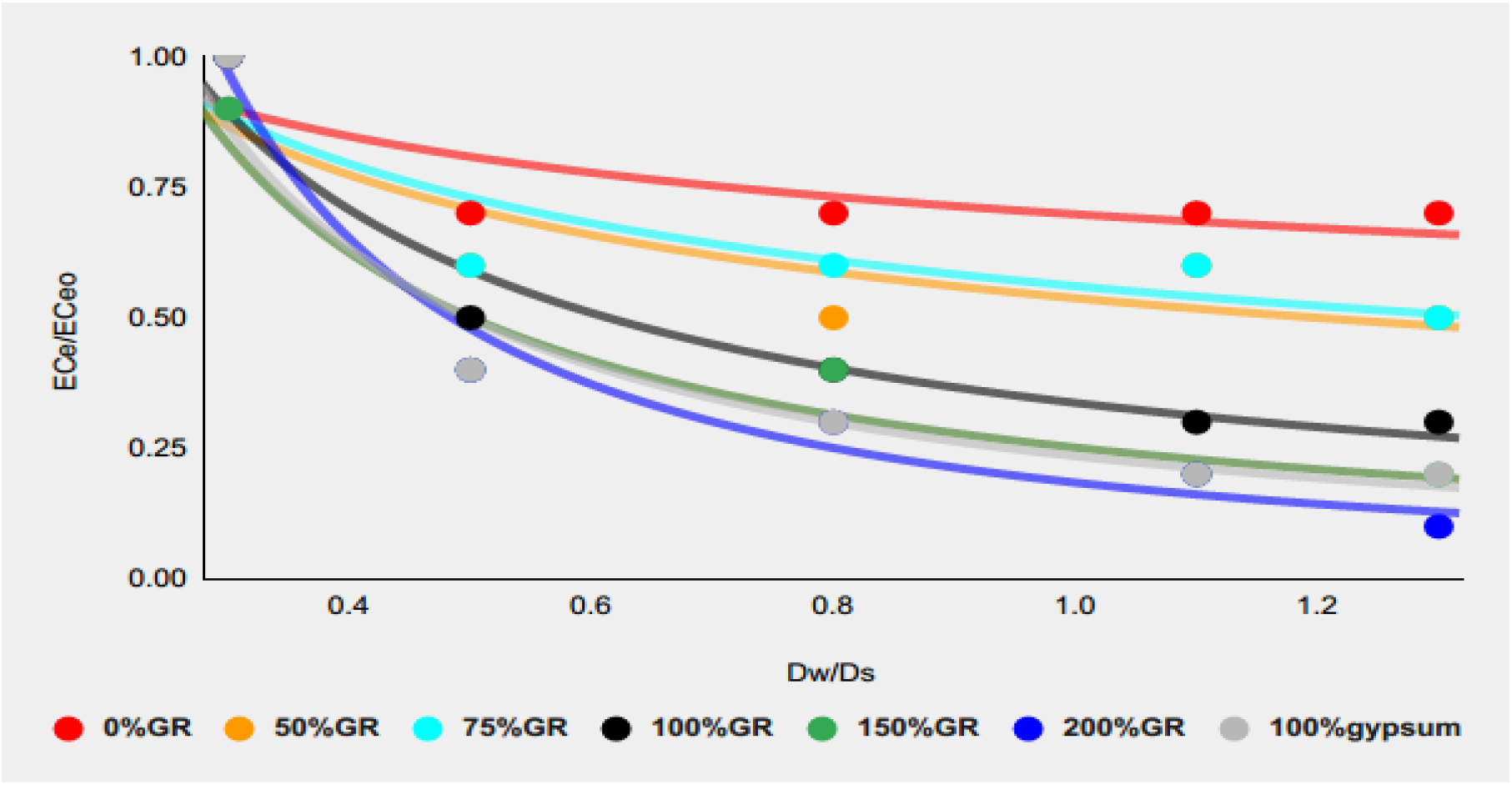
Desalinization leaching curve for saline-sodic soil amended with different PG rates

**Figure 5.**
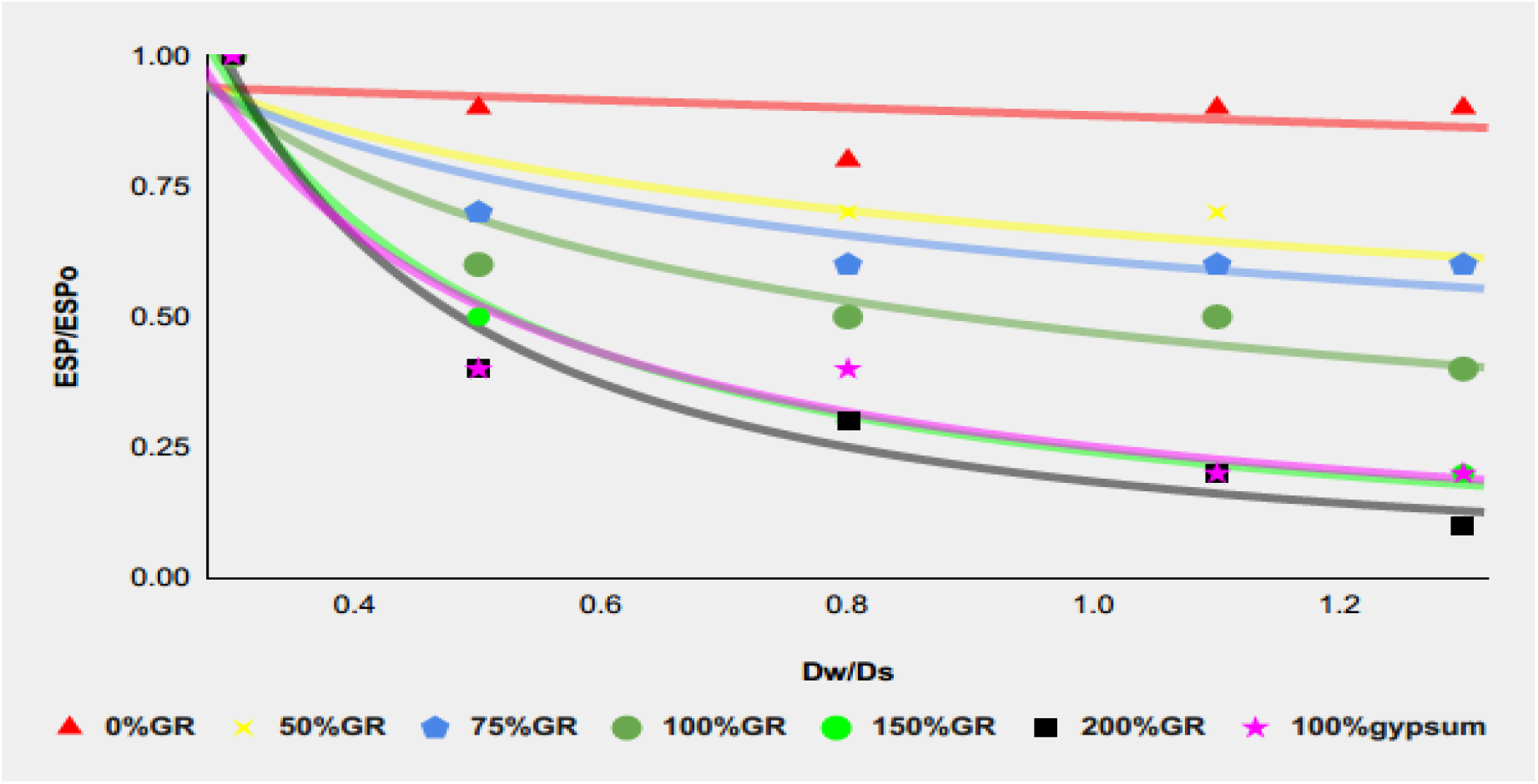
Desodification leaching curve of saline-sodic soil amended by different PG rates.

#### 3.3.1. Desalinization curve of the treated saline-sodic soil

The desalinization leaching curve (Figure 4) shows how the salinity of the soil changes as more water is leached through it. All PG treatment rates reduced the salinity of the soil, with the highest PG rate (200%GR) being the most effective compared to the natural gypsum and control treatments. The more water leached through the soil, the lower will be the salinity of the soil. Moreover. soils that were amended with higher PG rates required less water to reduce the salinity of the soil to an acceptable level (ECe < 4 dSm^−1^). This was achieved at PG application rates of 150%GR or higher and effective leaching depth (Dw/Ds) of 1.0 or less (< 4PV).

Therefore, following the equation developed for the desalinization curve, to remove 75% of the soluble salt from a soil with an initial salinity of 12 dS m^−1^ and bring it to ECe <4 dS m^−1^ for a soil depth of 20 cm, the required leaching water depth was 8.8 cm at a PG application rate of 150% GR, 66 cm at a PG application rate of 50% GR and 14.5 cm at application rate of 100% GR. This implies that higher PG rate is more effective than the lower rates and more effective than natural gypsum. [12] also reported similar trend that the efficiency of PG treatment (94.6%) was greater in reclaiming saline-sodic than that of natural gypsum (84.9%) at an application rate of 20.64 and 13.69 cm water respectively. Similar findings were also reported by [44] and the main reason for higher PG rates requiring less leaching water might be due to improvement of the permeability of water through the soil, making it easier for the soluble salts to be washed out. Additionally, according to [13] PG treatment improves the structure of the soil by improving the aggregation of soil particles. This is a common phenomenon when calcium ions (Ca^2+^) are added to the soil by using calcium bearing materials, and the improved soil structure leads to better hydraulic conductivity, which allows more effective leaching of soluble salts [44, 12].

#### 3.3.2 Desodification curve of treated saline-sodic soil

The desodification leaching curve (Figure 5) shows the changes in the ESP of the PG treated soil as more water is leached through it. Desodification leaching curves followed a similar trend with desalinization leaching curves, indicating that PG treatment is effective in reducing sodium from the exchange site (Figure 5). This confirms that addition of an amendment such as PG is required to reclaim saline-sodic soil besides leaching. The highest PG rate (200% GR) was the most effective in reducing sodicity though the safe level of ESP (ESP < 10) was achieved at combined application of PG at 150% GR or higher rates and an effective leaching depth (Dw/Ds) of 0.8 cm or less. This result agrees with the findings of a study by [44] that showed the efficiency of PG in reclaiming saline-sodic soils. This was influenced by the PG application rate and efficiency of leaching which influence the amount of calcium (Ca^2+^) released from the PG to replace the sodium (Na^+^).

Application of higher PG rates (100%GR, 150%GR and 200%GR) resulted in greater reductions in ESP and sodium content. This is because higher PG application rates provide more Ca^2+^ ions to displace the Na^+^ ions more than the natural gypsum. Similarly, [11] also found that application of higher PG rates provides more Ca^2+^ ions to displace the Na^+^ ions, which can also improve the efficiency of leaching.

### 3.4. Efficiency of PG and PV of leaching water on soil reclamation

The percentage of soluble salt and sodium removed from the soil (SSRE and SRE) for each PG application rate is shown in Table 5. The results showed that the efficiency of PG in removing sodium from saline-sodic soils was influenced by the PG application rate. Higher PG application rates (100%GR, 150%GR and 200%GR) resulted in greater reductions in ESP and sodium content while the efficiency of 100% natural gypsum is 69% which is lower than the above three PG rates. The highest PG application rate (200%GR) was the most effective at removing Na, with a removal rate of 75%, followed by the 150% GR and100%GR PG rates with a removal of 72% and 71% respectively. The control treatment was not effective at removing Na, and in fact, the SRE value was negative. This indicates that to remove Na from the soil, it is necessary to add a chemical amendment such as PG, and leach the soil with good quality water. Therefore, adding PG and subsequently leaching the soil are crucial steps in effective reclamation of saline-sodic soils. The present finding is consistent with the findings of [11], whereby the highest PG application rate (200% GR) was the most effective at removing Na, with a removal rate of 75%.

**Table 5.** Efficiency of PG amendment rates on reclamation of saline-sodic soils of Metehara Sugar Estate.

| PG rates | Soluble Salt Removal Efficiency (SSRE) | Na Removal Efficiency (SRE) |
| --- | --- | --- |
| 0%GR | 29% | -9% |
| 50%GR | 46% | 10% |
| 75%GR | 52% | 22% |
| 100%GR | 77% | 71% |
| 150%GR | 80% | 72% |
| 200%GR | 81% | 75% |
| 100% Gypsum | 75% | 69% |

The lowest K value (0.28) for desalinization was observed with 150% GR, followed by 200% GR (0.29) PG rate, while the highest K value (0.56) was observed with the control (Table 6).

**Table 6.** Efficiency of PG rates accompanied with PV of leaching water for saline-sodic soil of Metehara.

| PG rates<br>(Treatments) | Desalinization | Desodification |
| --- | --- | --- |
| | $K = (EC_e/EC_{eo}) \times (D_w/D_s)$ | $K = (ESP/ESP_o) \times (D_w/D_s)$ |
| 0%GR(control) | 0.56 | 0.98 |
| 50%GR | 0.51 | 0.80 |
| 75%GR | 0.50 | 0.70 |
| 100%GR | 0.32 | 0.26 |
| 150%GR | 0.28 | 0.25 |
| 200%GR | 0.29 | 0.22 |
| 100% Gypsum | 0.33 | 0.27 |

This implies that less water is required to leach soil treated with the highest PG rate when compared with the control and 100% natural gypsum. Similarly, the lowest K value (0.22) for desodification was observed with the highest PG rate (200% GR), indicating that less water is required with the highest PG rate. This may be due to the improvement of soil structure by treatment and enhancing removal of more Na and its effects, indicating that higher PG rates are more effective in doing so. The fact that the lowest K value for desodification was observed with the highest PG rate is also consistent with the findings of [48], and [24]. Also, [12] and [43] reported application of PG at high rate needed less leaching water. This is because PG helps removal of more Na from the soil, and improves the structure of the soil making it easier for water to flow through.

According to the current findings, both desalinization and desodification were more effective with PG application rates of (≥100%GR) than 100% pure gypsum and control. [13] also reported similar finding in that higher PG rate had high ability to reduce salts compared to natural gypsum.

### 3.5. Conclusions

The results of the column experiment that application PG combined with leaching water of different quantities improved most salinity and sodicity-related soil properties. Using PG at a rate of 100% GR or higher (≥13 tons PG per hectare) with the first three rounds of leaching water was the most effective way to reduce soil salinity and sodium content of the saline-sodic soils of Metehara Sugar Estate. This was better than using lower amounts of PG (<100% GR) or natural gypsum (100% G) alone. Therefore, it is necessary to apply at least 13 tons of PG per hectare together with 3 to 4 pore volumes of leaching water to achieve the safe level of salinity (ECe < 4 dSm^−1^) and sodicity (ESP < 10%) for most crops. Overall, the results suggest that PG is a promising amendment for reclaiming saline-sodic soils and improving soil health.

While the laboratory experiments showed promising results, additional research is needed in greenhouse and field settings to confirm the present findings in natural environments. This will help better understand the potential of PG as a resource for reclaiming salt-affected soils.

## Acknowledgements

The authors are grateful to the Metehara Sugar state farm field staff and Haramaya University laboratory technician who support us during field soil sample collection and laboratory analysis.

## Author contributions

**Conceptualization:** Kebenu Feyisa, Kibebew Kibret, Sheleme Beyene, Lemma Wogi

**Data curation:** Kebenu Feyisa, Kibebew Kibret, Sheleme Beyene, Lemma Wogi

**Formal analysis:** Kebenu Feyisa, Kibebew Kibret, Sheleme Beyene, Lemma Wogi

**Funding acquisition:** Kebenu Feyisa, Kibebew Kibret, Sheleme Beyene

**Investigation:** Kebenu Feyisa, Kibebew Kibret, Sheleme Beyene, Lemma Wogi

**Methodology:** Kebenu Feyisa, Kibebew Kibret, Sheleme Beyene, Lemma Wogi

**Project administration:** Kebenu Feyisa, Kibebew Kibret, Sheleme Beyene, Lemma Wogi

**Resources:** Kebenu Feyisa, Kibebew Kibret, Sheleme Beyene, Lemma Wogi

**Software:** Kebenu Feyisa, Kibebew Kibret, Sheleme Beyene, Lemma Wogi

**Supervision:** Kibebew Kibret, Sheleme Beyene, Lemma Wogi

**Validation:** Kebenu Feyisa, Kibebew Kibret, Sheleme Beyene, Lemma Wogi

**Visualization:** Kebenu Feyisa, Kibebew Kibret, Sheleme Beyene, Lemma Wogi

**Writing-original draft:** Kebenu Feyisa

**Writing-review and editing:** Kebenu Feyisa, Kibebew Kibret, Sheleme Beyene, Lemma Wogi

## Data availability statement

All relevant data are within the manuscript in tables and figures.

